# Species, causes, and outcomes of wildlife rehabilitation in New York State

**DOI:** 10.1101/860197

**Authors:** Melissa Hanson, Nicholas Hollingshead, Krysten Schuler, William F. Siemer, Patrick Martin, Elizabeth M. Bunting

## Abstract

Wildlife rehabilitation is a publicly popular though highly controversial practice. State wildlife agencies frequently debate the ecological impact of rehabilitation. Analysis of case records could inform that debate by clarifying and quantifying the causes for rehabilitation, species involved, and treatment outcomes. This information could aid in the ability of regulatory agencies and rehabilitators to make informed decisions and gain insight into causes of species decline. In New York, rehabilitators are licensed by the Department of Environmental Conservation (NYSDEC) and thus, are required to submit annual reports. Between 2012-2014, 59,370 individual wildlife cases were seen by licensed rehabilitators comprising 31,229 (52.6%) birds, 25,490 (42.9%) mammals, 2,423 (4.1%) reptiles, and 73 (0.1%) amphibians. We identified patterns among taxonomic representation, reasons for presentation, and disposition. Major causes of presentation were trauma (n = 22,672, 38.2%) and orphaning (n = 21,876, 36.8%), with habitat loss (n =3,746, 6.3%), infectious disease (n = 1,992, 3.4%), and poisoning or toxin exposure (n = 864, 1.5%) playing lesser roles. The overall release rate for animals receiving care was 50.2%; 45.4% were either euthanized or died during the rehabilitation process. A relatively small number (0.3%) were permanently non-releasable and placed in captivity, and 4.1% had unknown outcomes. In comparison to data from 1989, wildlife submissions have increased (annual mean 12,583 vs 19,790), as has the release rate, from 44.4% to 50.2%. Utilizing a large data set allowed us to fill knowledge gaps, which can help inform management by both the rehabilitators and the state agencies that regulate them, deepening understanding of the scope and impacts of wildlife rehabilitation.

## Introduction

Ethical questions and skepticism over the ecological benefits have fueled debate on rehabilitative treatment of wild animals [1,2]. The value of rehabilitation for individual animals is also controversial, with little knowledge of release rates, and some arguing that stressors placed on animals undergoing care at the rehabilitation facilities may be as traumatizing to the animal as the inciting event [3–5]. In contrast, advocates urge that the need for rehabilitation often arises from anthropogenic causes, and humans therefore have a moral obligation to rectify their impact [6]. In addition, rehabilitation provides people with close contact with wildlife, potentially increasing knowledge of wild species and factors contributing to their declines, which can have positive impacts on local biodiversity conservation [6].

Without basic quantifiable information, this debate has been left at a standstill. Here, we report large-scale epidemiologic data to provide a foundation for understanding the scope and potential impacts of wildlife rehabilitation.

Wildlife rehabilitation is the practice of caring for sick, injured, orphaned, and displaced wild animals with the primary goal of returning them to their natural habitat. Wildlife rehabilitators, typically dedicated private citizens or small non-profit centers, may invest significant time and personal resources in this activity: the National Wildlife Rehabilitators Association (NWRA) reports that a rehabilitator volunteers an average of 32-36 hours per week through the spring and summer of each calendar year [7]. The roles of a rehabilitator are multi-disciplinary—from public outreach, education, and advocacy to husbandry, nursing, and collaboration with veterinary professionals. The rehabilitator plays a primary role in the ultimate disposition of the animal in question, from selecting an appropriate release location to consulting on the euthanasia of individuals that cannot be rehabilitated.

Of equal importance is the responsibility rehabilitators hold to collect and report information about the animals they care for—thus contributing to the understanding of local and national populations, as well as the overall health and well-being of the habitat shared between wildlife and people [8]. In particular, global declines in wildlife numbers raise important questions regarding specific causes of these trends. Recent studies estimate that one third of terrestrial wildlife admissions to rehabilitation facilities and 70% of loggerhead sea turtle [*Caretta caretta*] stranding events that required rehabilitative intervention were attributed to anthropogenic activity [1,9].

Part of the challenge with resolving conflicts, answering questions about impacts, or even improving basic treatment strategies is that there are only limited data on wildlife rehabilitation to support informed decisions. Studies are limited due to the dispersed structure of the rehabilitation community and the lack of standardized digital records.

In 1980, the New York State Department of Environmental Conservation (DEC) began issuing wildlife rehabilitation licenses to qualified private citizens, and since 1985 the agency has required them to submit annual reports on paper forms. Therefore, data regarding rehabilitation case numbers, species affected, causes of presentation, and treatment outcomes/disposition was collected, but not easily accessible. Following digitization, we report large-scale epidemiological evaluation of 59,370 individual animal records submitted from wildlife rehabilitators across New York State (NYS) over three years: 2012-2014. Previous analyses were available from 1989, so further comparison is possible to detect changes in the system over time.

This large-scale analysis of wildlife rehabilitation case data may assist rehabilitators in allocating limited treatment resources and improving care. State wildlife agencies tasked with oversight can benefit by improving their understanding of the scope, needs, and impact of rehabilitation. Closer examination of several species of interest also suggests the potential for population impacts, further informing the debate regarding the scope and utility of wildlife rehabilitation.

## Materials and methods

Licensed wildlife rehabilitators in New York are required to record case information on a weekly basis using a paper Wildlife Rehabilitator Log (WRL) form provided by the DEC. Annual summaries and all WRL forms are submitted to the agency by December 1 of each calendar year. Data on these handwritten logs from the three reporting years ending December 1, 2012, 2013, and 2014 were manually transcribed into digital spreadsheets, and records from all years were then combined into a single Microsoft Office Access® database (Microsoft Corporation, Redmond, Washington, USA, 90852). All database records were reviewed for accuracy and formatting consistency. Simple data entry errors by the wildlife rehabilitators (such as minor incorrect spelling of species or place names) were corrected. Species names were standardized using currently accepted taxonomic lists [10,11]. Terms used for causes of presentation/distress and dispositions were taken from the DEC classifications on the forms. Incomplete and inaccurate records, as well as records for domestic and captive animals, were excluded from the analysis.

Species reported were aggregated into four primary groupings and 17 secondary groupings commonly used by rehabilitators as follows: birds (Columbiformes, raptors, passerines, waterfowl, and other), mammals (large mammals, small herbivores, and small carnivores), reptiles (turtles, snakes, lizards), rabies vector species (bats, raccoons [*Procyon lator*], and striped skunk [*Mephitus mephitis*]), amphibians (frogs and salamanders) and unknown. The number of reptiles other than turtles were too few for meaningful analysis, so this category was reduced to turtles alone. A list of species included in the analysis for each grouping is provided in Tables 2–10.

On the WRL form, the causes of distress included 15 primary categories and 42 subcategories. For analytical purposes, the 15 primary categories were aggregated into six groupings: 1) orphaned (orphaned, orphaned due to uneccessary human intervention, developmental abnormality); 2) trauma (accidental entrapment, collision, entanglement, injured by other animal or human, mechanical injury due to gun/arrow mower or trap); 3) infectious (bacterial infection, parasitism, viral disease); 4) poisoning/toxin (poison or toxin ingestion, soaked or similar damage); 5) habitat loss (human disturbance, i.e. tree-cutting, building construction, and natural disturbance, i.e. flood, fire, etc); and 6) unknown. Because of the great interest in understanding the impact of domestic cats on declining wildlife numbers [12], trauma from cat attacks have been disaggregated and presented where appropriate. A small subset of records had more than one distress code provided. For these cases, each combination was evaluated and one reported cause was selected as the primary reason for presentation to a rehabilitation facility. For example, all causes of trauma or poisoning/toxin were selected as primary when combined with orphaning, while parasitism was considered secondary to orphaning.

The WRL disposition categories included: 1) died under/prior to care; 2) euthanized; 3) permanently non-releasable and transferred to a person with a valid NYS license; 4) permanently non-releasable and transferred to an educational institute with a valid NYS license; 5) released; 6) still under care; 7) transferred to another licensed wildlife rehabilitator; and 8) unknown. When cases were marked as being transferred to a second rehabilitator, only the secondary record was included in the analysis in order to avoid duplication. The disposition categories were combined into the following groups for analysis: 1) died (died under/prior to care, euthanized); 2) released; 3) non-releasable (placed under permanent care of licensed individual or educational institution); and 4) unknown (still under care or unknown).

We divided release rates by species and causes of distress. Cases and release rates were reported for the most common species within each species grouping. Individual species of interest in New York, including bald eagles (*Haliaeetus leucocephalus*), eastern box turtles (*Terrapene carolina carolina*), and wood ducks (*Aix sponsa*), were examined to provide information on the scope of rehabilitation efforts relative to population estimates derived from the number of resident breeding pairs.

Because licensed wildlife rehabilitators in the State of New York are required to submit annual case logs, the 2012-2014 dataset was effectively a complete census of all animals under the care of wildlife rehabilitators during that three-year time period. Our basic summary metrics for this census include proportions and counts and negate the need for inferential statistics. However, to analyze differences in rehabilitation category counts between different time periods, we conducted a chi-square goodness-of-fit test to ascertain whether category counts in the recent past (2012-2014) were statistically different from category counts collected in 1989 [13]. We performed the statistical analyses using Prism 7 (GraphPad, San Diego, California, USA, 92108).

## Results

Between 2012 and 2014, rehabilitators reported 59,370 wildlife rehabilitation cases to the DEC. The cases were primarily birds (n=31,229, 52.6%) and mammals (n=25,490, 42.9%) with lesser numbers of reptiles (n=2,423, 4.1%) and amphibians (n=73, 0.1%). The remainder (n=155, 0.3%) were not clearly specified.

We evaluated intake records from 703 uniquely identifiable rehabilitators. Each year, between 450 and 457 of these rehabiltiators had at least one case and submitted their WRL. Individual wildlife rehabilitators could be identified for 53,328 cases (89.8%). The majority of rehabilitators (74.9%) saw 25 or fewer animals per annum, accounting for 7,990 (15.0%) cases. By comparison, 21,925 animals (41.1%) were seen by the 2.3% of rehabilitators who saw more than 300 animals annually (Table 1).

**Table 1.**
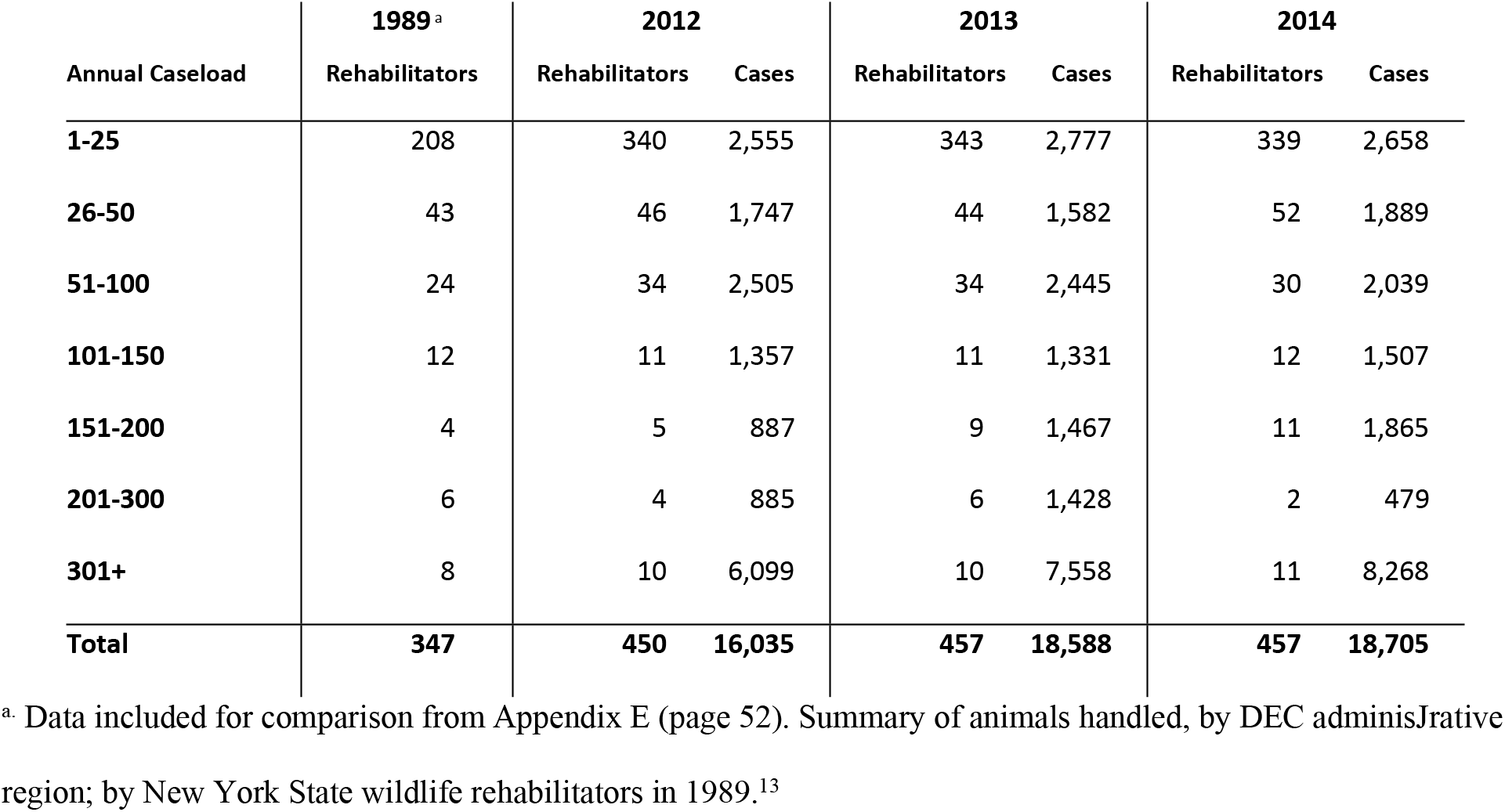
Annual caseload reported by individual wildlife rehabilitators in New York 2012-2014.

A cause of distress was provided for 86.2% of the records, with 2.4% (n = 1,442) of the cases having more than one cause of distress code given. Of those cases with multiple codes, 113 had codes within the same primary distress category and were not subject to comparison for purposes of analysis. Of the remaining cases, 1,061 had causes of distress in two primary categories, and 187 in more than two primary categories.

When the cases were aggregated into the six general groupings, trauma was the most common overall cause of submission (n=22,674, 38.2%) followed by orphaning (n= 21,876, 36.8%); habitat loss (n=3,746 6.3%); infectious disease n=1,992, 3.4%); and poisoning/toxin (n=864, 1.5%). The remaining 8,220 (13.8%) cases had an unknown cause of distress or did not provide a cause of distress.

Final dispositions were reported for 95.7% (n=56,921) of cases. Approximately half of the animals presented (n=29,793, 50.2%) were returned to the wild at the end of their rehabilitation. During the study period, the release rate was greater for animals that remained under care for more than 2 days (70.8%; 95% CI: 70.1-71.5%) than it was for animals in care for 2 days or less (15.7%; 95% CI: 15.1-16.4%) [*χ*^2^ (1 df, N=28,054) = 8299, 1, p < 0.00001]. The overall release rate in the 2012-2014 time period improved significantly from 44.4% in 1989 [*χ*^2^ (1 df, N=71,370) = 137.7, p<0.00001] [13].

Rehabilitators reported that 26,957 animals (45.4%) either died naturally or were euthanized during rehabilitation. Other dispositions were rare in comparison; 1,967 animals (3.3%) remained under the care of the rehabilitator at the time of reporting, and 171 animals (0.3%) were deemed permanently non-releasable. The non-releasable animals were then moved to an educational facility or placed under the care of a licensed professional at another location. Final dispositions were not available for 482 cases (0.8%).

Waterfowl (Table 2) had the highest overall rate of release (66.4%) when compared with other groups. Of the 4,119 waterfowl that were admitted, 2,718 were eventually released. Of these successfully released waterfowl, 1,441 individuals had arrived at the facilities due to orphaning, while 606 had arrived as a result of trauma. The mallard duck [*Anas platyrhynchos*] had the highest overall rehabilitation success rate of all species evaluated with 77.8% being released (2,033 admitted; 1,582 released). For comparison, 224 (50.4%) of the 444 admitted wood ducks were successfully released (Table 2). Wood ducks represented 10.7% of the waterfowl presented to wildlife rehabilitators and possessed the lowest release rate amongst the most commonly seen waterfowl species. Most (84%) presented as orphans, and had substantially lower success rates compared to other orphaned waterfowl (52% vs. 78% for all waterfowl).

**Table 2.** Cause of presentation and corresponding disposition for waterfowl in rehabilitation.

|  | Habitat Loss |  | Infectious |  | Orphaned |  | Poisoning/Toxin |  | Trauma |  | Unknown |  | Total |  |
| --- | --- | --- | --- | --- | --- | --- | --- | --- | --- | --- | --- | --- | --- | --- |
|  | Cases | % Rel.<sup>a</sup> | Cases | % Rel. | Cases | % Rel. | Cases | % Rel. | Cases | % Rel. | Cases | % Rel. | Cases | % Rel. |
| **Mallard** | 32 | 66% | 10 | 60% | 1,138 | 85% | 11 | 91% | 528 | 66% | 314 | 73% | 2,033 | 78% |
| **Canada Goose** | 9 | 56% | 19 | 68% | 245 | 87% | 19 | 26% | 417 | 42% | 200 | 56% | 909 | 58% |
| **Diving Ducks<sup>b</sup>** | 140 | 61% | 16 | 56% | 75 | 61% | 33 | 48% | 101 | 46% | 169 | 57% | 534 | 56% |
| **Wood Duck** | 3 | 100% | 0 | 0% | 374 | 52% | 0 | 0% | 38 | 29% | 29 | 55% | 444 | 50% |
| **Other<sup>c</sup>** | 12 | 58% | 8 | 38% | 26 | 77% | 4 | 75% | 87 | 28% | 62 | 50% | 199 | 44% |
| **Total** | **196** | **62%** | **53** | **58%** | **1,858** | **78%** | **67** | **51%** | **1,171** | **52%** | **774** | **63%** | **4,119** | **66%** |

<sup>a</sup>. % Rel= Percent Released
<sup>b</sup>. Includes: Black scoter [*Melanitta americana*], Canvasback [*Aythya valisineria*], Common merganser [*Mergus merganser*], Greater scaup [*Aythya marila*], Hooded merganser [*Lophodytes cucullatus*], Lesser scaup [*Aythya affinis*], Mute swan [*Cygnus olor*], Red-breasted merganser [*Mergus serrator*], Redhead [*Aythya americana*], Ring-necked duck [*Aythya collaris*], Surf scoter [*Melanitta perspicillata*], and White-winged scoter [*Melanitta deglandi*]
<sup>c</sup>. Includes: American black duck [*Anas rubripes*], American wigeon [*Mareca americana*], Brant [*Branta bernicla*], Bufflehead [*Bucephala albeola*], Common eider [*Somateria mollissima*], Common goldeneye [*Bucephala clangula*],
gadwall [*Mareca strepera*], Long-tailed duck [*Clangula hyemalis*], Northern shoveler [*Spatula clypeata*], Ruddy duck [*Oxyura jamaicensis*], Snow goose [*Anser caerulescens*], Trumpeter swan [*Cygnus buccinator*], and Tundra swan [*Cygnus columbianus*]

Raptors (Table 3) had the lowest collective release rate (n = 1,465/3,232, 45.3%), though release success varied greatly depending on reason for admittance; orphaning had an 84.3% release rate while trauma had a 40.4% release rate. This difference in disposition based on presenting distress was consistent among raptor species, including the commonly presenting red-tailed hawk [*Buteo jamaicensus*], which was represented by 912 cases. Trauma was the most common cause of distress for raptor species (57.1%), followed by unknown/unspecified causes (20.2%), and orphaning (12.2%).

**Table 3.** Cause of presentation and corresponding disposition for raptors in rehabilitation.

|  | Habitat Loss |  | Infectious |  | Orphaned |  | Poisoning/Toxin |  | Trauma |  | Unknown |  | Total |  |
| --- | --- | --- | --- | --- | --- | --- | --- | --- | --- | --- | --- | --- | --- | --- |
|  | Cases | % Rel.<sup>a</sup> | Cases | % Rel. | Cases | % Rel. | Cases | % Rel. | Cases | % Rel. | Cases | % Rel. | Cases | % Rel. |
| **Red-tailed Hawk** | 23 | 70% | 75 | 23% | 34 | 76% | 21 | 19% | 548 | 36% | 211 | 34% | 912 | 37% |
| **Eastern Screech-Owl** | 32 | 84% | 7 | 71% | 113 | 84% | 0 | 0% | 267 | 47% | 73 | 55% | 492 | 60% |
| **American Kestrel** | 16 | 94% | 1 | 100% | 174 | 91% | 2 | 50% | 95 | 53% | 42 | 50% | 330 | 75% |
| **Barred Owl** | 8 | 38% | 4 | 0% | 16 | 81% | 3 | 100% | 257 | 51% | 30 | 27% | 318 | 50% |
| **Great Horned Owl** | 14 | 64% | 19 | 26% | 20 | 65% | 7 | 29% | 140 | 38% | 73 | 37% | 273 | 40% |
| **Cooper's Hawk** | 7 | 71% | 18 | 11% | 8 | 100% | 7 | 43% | 181 | 37% | 49 | 29% | 270 | 37% |
| **Broad-winged Hawk** | 5 | 80% | 5 | 20% | 8 | 63% | 1 | 0% | 60 | 38% | 13 | 38% | 92 | 41% |
| **Sharp-shinned Hawk** | 1 | 100% | 4 | 25% | 0 | 0% | 0 | 0% | 56 | 38% | 18 | 17% | 79 | 33% |
| **Bald Eagle** | 5 | 80% | 5 | 20% | 3 | 67% | 7 | 43% | 14 | 36% | 21 | 48% | 55 | 45% |
| **Other<sup>b</sup>** | 15 | 27% | 14 | 7% | 20 | 65% | 11 | 18% | 228 | 31% | 123 | 37% | 411 | 33% |
| **Total** | **126** | **70%** | **152** | **22%** | **396** | **84%** | **59** | **31%** | **1,846** | **40%** | **653** | **38%** | **3,232** | **45%** |

<sup>a</sup> % Rel= Percent Released
<sup>b</sup> Includes: Barn owl [*Tyto alba*], Black vulture [*Coragyps atratus*], Golden eagle [*Aquila chrysaetos*], Long-eared owl [*Asio otus*], Merlin [*Falco columbarius*], Northern goshawk [*Accipiter gentilis*], Northern harrier [*Circus*
*hudsonius*], Northern saw-whet owl [*Aegolius acadicus*], Osprey [*Pandion haliaetus*], Peregrine falcon [*Falco* *peregrinus*], Red-shouldered hawk [*Buteo lineatus*], Rough-legged hawk [*Buteo lagopus*], Short-eared owl [*Asio* *flammeus*], Snowy owl [*Bubo scandiacus*], and Turkey vulture [*Cathartes aura*].

Bald eagles represented less than 2% of raptors presented to wildlife rehabilitation centers. Approximately one-quarter of all admissions were a result of trauma, with a release rate of 36% closely mirroring that for other trauma-victim raptors. The next most common known cause for presentation of bald eagles was poisoning/toxin (12.7%). Bald eagles presented more commonly with this distress than any other raptor species included in this analysis, but only 3/7 (43%) individuals were successfully rehabilitated.

Passerines (Table 4) were the second most likely group to be submitted for rehabilitation (13,100) and had an overall release rate of 46.1% (n = 6,035/13,100). The American crow [*Corvus brachyrhynchos*] had the lowest overall release rate for any species evaluated in this study (n = 142/544, 26.1%). According to the rehabilitators’ reports, crows presented to wildlife centers primarily for infectious disease 14.5% of the time (n = 79/544), far more frequently than all other passerines (2%). These crows had lower release rates (3.8%) than all other passerines with infectious disease (n = 77/254, 30.3%). Overall, crows had the lowest release rates for every category of distress, except orphaning.

**Table 4.**
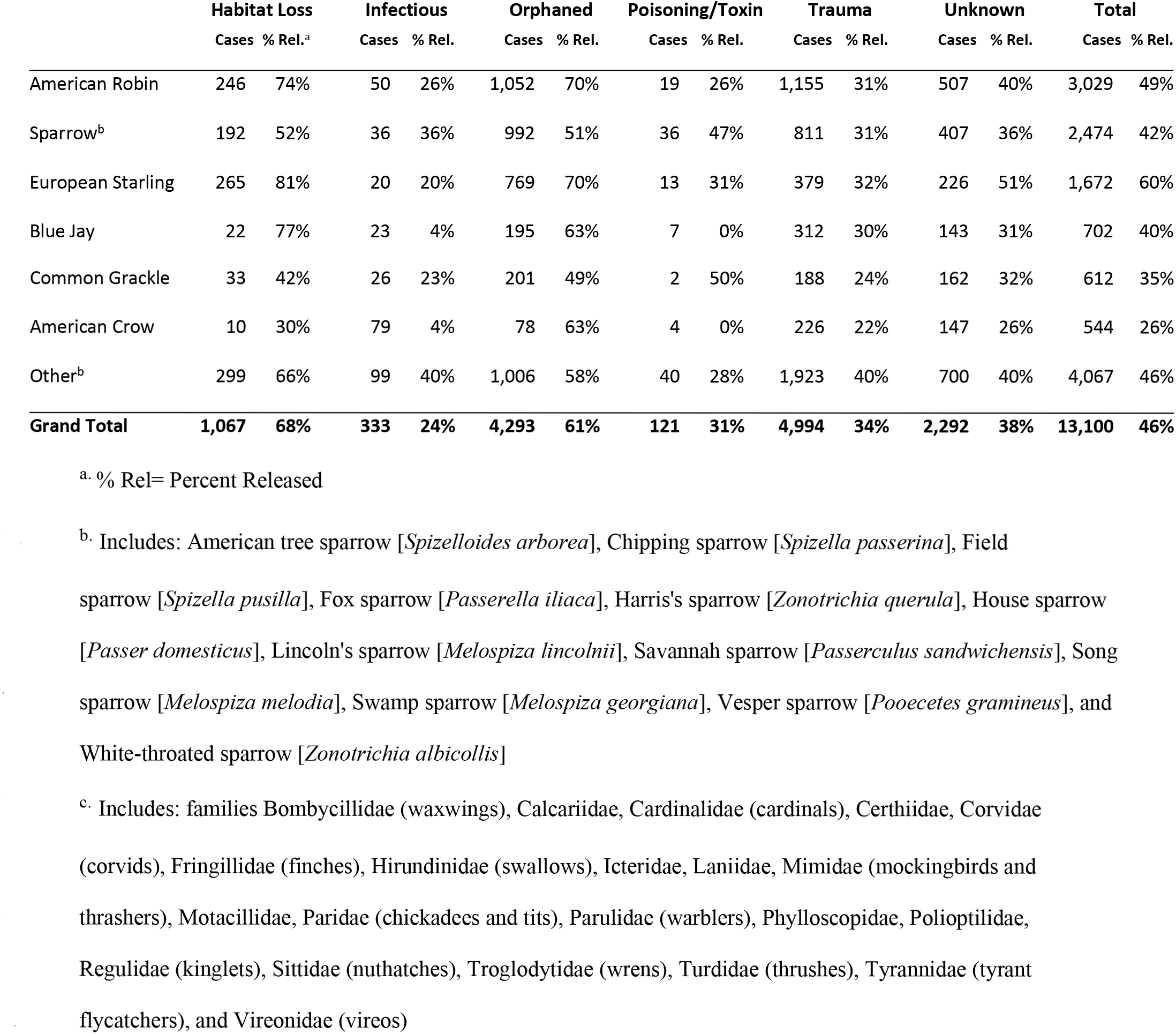
Cause of presentation and corresponding disposition for passerines in rehabilitation.

|  | Habitat Loss |  | Infectious |  | Orphaned |  | Poisoning/Toxin |  | Trauma |  | Unknown |  | Total |  |
| --- | --- | --- | --- | --- | --- | --- | --- | --- | --- | --- | --- | --- | --- | --- |
|  | Cases | % Rel.<sup>a</sup> | Cases | % Rel. | Cases | % Rel. | Cases | % Rel. | Cases | % Rel. | Cases | % Rel. | Cases | % Rel. |
| American Robin | 246 | 74% | 50 | 26% | 1,052 | 70% | 19 | 26% | 1,155 | 31% | 507 | 40% | 3,029 | 49% |
| Sparrow<sup>b</sup> | 192 | 52% | 36 | 36% | 992 | 51% | 36 | 47% | 811 | 31% | 407 | 36% | 2,474 | 42% |
| European Starling | 265 | 81% | 20 | 20% | 769 | 70% | 13 | 31% | 379 | 32% | 226 | 51% | 1,672 | 60% |
| Blue Jay | 22 | 77% | 23 | 4% | 195 | 63% | 7 | 0% | 312 | 30% | 143 | 31% | 702 | 40% |
| Common Grackle | 33 | 42% | 26 | 23% | 201 | 49% | 2 | 50% | 188 | 24% | 162 | 32% | 612 | 35% |
| American Crow | 10 | 30% | 79 | 4% | 78 | 63% | 4 | 0% | 226 | 22% | 147 | 26% | 544 | 26% |
| Other<sup>b</sup> | 299 | 66% | 99 | 40% | 1,006 | 58% | 40 | 28% | 1,923 | 40% | 700 | 40% | 4,067 | 46% |
| **Grand Total** | **1,067** | **68%** | **333** | **24%** | **4,293** | **61%** | **121** | **31%** | **4,994** | **34%** | **2,292** | **38%** | **13,100** | **46%** |

<sup>a</sup>. % Rel= Percent Released
<sup>b</sup>. Includes: American tree sparrow [*Spizelloides arborea*], Chipping sparrow [*Spizella passerina*], Field sparrow [*Spizella pusilla*], Fox sparrow [*Passerella iliaca*], Harris's sparrow [*Zonotrichia querula*], House sparrow [*Passer domesticus*], Lincoln's sparrow [*Melospiza lincolni*], Savannah sparrow [*Passerculus sandwichensis*], Song sparrow [*Melospiza melodia*], Swamp sparrow [*Melospiza georgiana*], Vesper sparrow [*Pooecetes gramineus*], and White-throated sparrow [*Zonotrichia albicollis*]
<sup>c</sup>. Includes: families Bombycillidae (waxwings), Calcariidae, Cardinalidae (cardinals), Certhiidae, Corvidae (corvids), Fringillidae (finches), Hirundinidae (swallows), Icteridae, Laniidae, Mimidae (mockingbirds and thrashers), Motacillidae, Paridae (chickadees and tits), Parulidae (warblers), Phylloscopidae, Polioptilidae, Regulidae (kinglets), Sittidae (nuthatches), Troglodytidae (wrens), Turdidae (thrushes), Tyrannidae (tyrant flycatchers), and Vireonidae (vireos)

The Columbiforme group (Table 5) was dominated by the pigeon [*Columba* spp.], which was the most frequently treated bird in the study. Trauma was the most common cause of admittance for both Columbiformes and passerines, and the Pigeon had the highest release rates for trauma victims in both of these groups (n = 1,006/2,253, 44.7%).

**Table 5.** Cause of presentation and corresponding disposition for pigeons and doves in rehabilitation.

|  | Habitat Loss |  | Infectious |  | Orphaned |  | Poisoning/Toxin |  | Trauma |  | Unknown |  | Total |  |
| --- | --- | --- | --- | --- | --- | --- | --- | --- | --- | --- | --- | --- | --- | --- |
|  | Cases | % Rel.<sup>a</sup> | Cases | % Rel. | Cases | % Rel. | Cases | % Rel. | Cases | % Rel. | Cases | % Rel. | Cases | % Rel. |
| **Pigeon<sup>b</sup>** | 120 | 68% | 900 | 38% | 885 | 66% | 407 | 48% | 2,253 | 45% | 1,005 | 39% | 5,570 | 47% |
| **Dove<sup>c</sup>** | 37 | 81% | 38 | 21% | 259 | 68% | 15 | 13% | 804 | 32% | 292 | 46% | 1,445 | 42% |
| **Total** | **157** | **71%** | **938** | **37%** | **1,144** | **66%** | **422** | **46%** | **3,057** | **41%** | **1,297** | **40%** | **7,015** | **46%** |

<sup>a</sup>. % Rel= Percent Released
<sup>b</sup>. Includes: Rock pigeon [*Columba livia*]
<sup>c</sup>. Includes: Mourning dove [*Zenaida macroura*] and Ring-necked dove [*Streptopelia capicola*]

The large mammal group (Table 6) was the smallest group submitted to rehabilitation. Approximately half of all admissions were orphaned, and 85.9% (n = 1,441/1,677) of the large mammals treated were neonates and juveniles, the highest proportion of all groups. Though trauma was a less common reason for admittance, large mammal trauma patients had a relatively low release rate (n = 165/562, 29.4%).

**Table 6.**
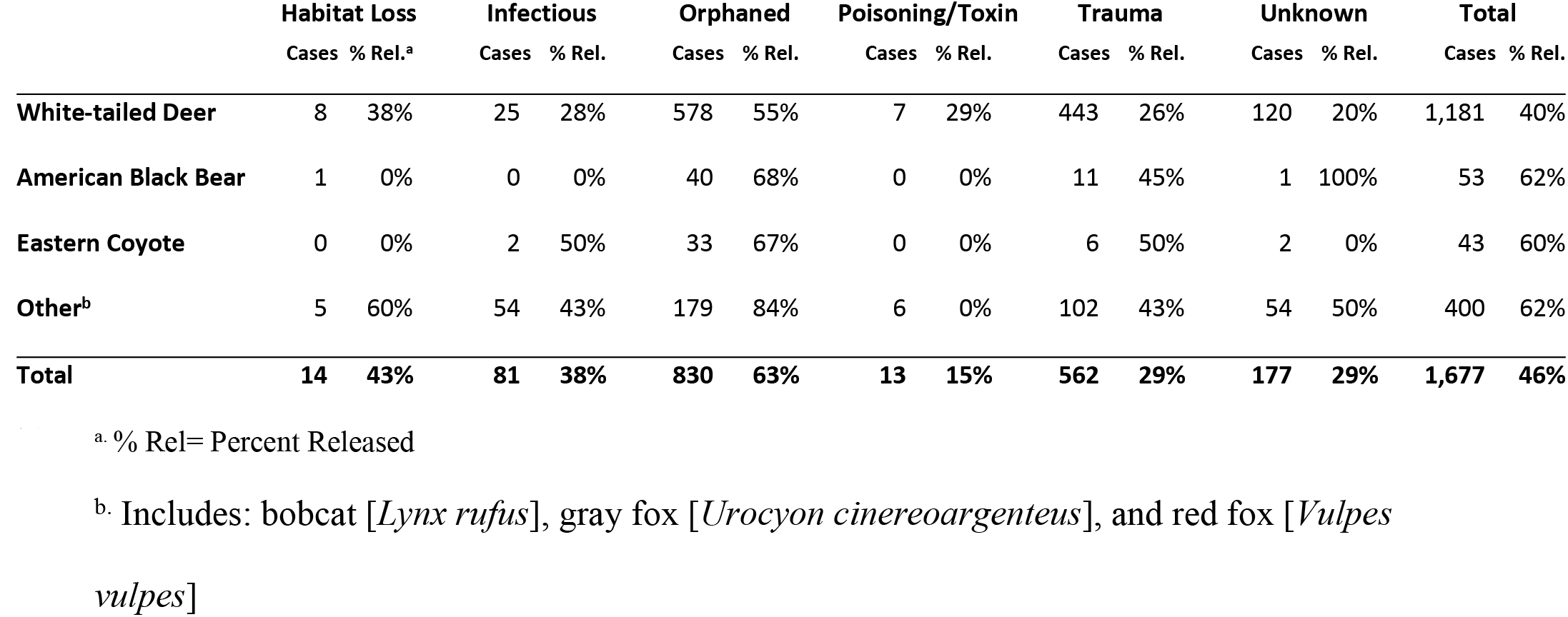
Cause of presentation and corresponding disposition for large mammals in rehabilitation.

The small herbivores (Table 7) represented the group most likely to be submitted for rehabilitation (18,629) and a large proportion (n = 16,282/18,629, 87.4%) were either eastern cottontails [*Sylvilagus floridanus*] or eastern grey squirrels [*Sciurius carolinensis*]. The overall release rates for grey squirrels and eastern cottontails were 65.4% (n = 4,876/7,459), and 44.7% (n = 3,942/8,823), respectively. The leading primary cause of distress for eastern cottontails was trauma (n = 4,452/8,823, 50.5%), which was typically associated with a poor outcome, indicated by a release rate of just 33.7% (n = 1,500/4,452). More specifically, 2,807 rabbits (31.8% of all admissions for the species) were presented to wildlife rehabilitators for trauma associated with a domestic dog or cat attacks. Trauma was much less common in grey squirrels (n =1,367/7,459, 18.3%), whereas orphaning was the leading cause of distress (n = 4,452/7,459, 59.7%). Orphaning was also frequent for eastern cottontails, as the second most common reason for admittance (n = 3,452/8,823, 39.1%). The release rate for orphaned squirrels was higher (n = 3,340/4,452, 75.0%) than that for eastern cottontails (n = 2,011/3,452, 58.3%) (Table 7).

**Table 7.** Cause of presentation and corresponding disposition for small herbivores in rehabilitation.

|  | Habitat Loss |  | Infectious |  | Orphaned |  | Poisoning/Toxin |  | Trauma |  | Unknown |  | Total |  |
| --- | --- | --- | --- | --- | --- | --- | --- | --- | --- | --- | --- | --- | --- | --- |
|  | Cases | % Rel.<sup>a</sup> | Cases | % Rel. | Cases | % Rel. | Cases | % Rel. | Cases | % Rel. | Cases | % Rel. | Cases | % Rel. |
| Eastern Cottontail | 437 | 54% | 27 | 15% | 3,452 | 58% | 36 | 31% | 4,452 | 34% | 419 | 43% | 8,823 | 45% |
| East. Gray Squirrel | 942 | 74% | 92 | 26% | 4,452 | 75% | 37 | 43% | 1,367 | 39% | 569 | 46% | 7,459 | 65% |
| Other Squirrel Sp.<sup>b</sup> | 119 | 61% | 3 | 33% | 363 | 66% | 2 | 0% | 165 | 59% | 57 | 61% | 709 | 63% |
| Woodchuck | 3 | 67% | 9 | 33% | 191 | 82% | 4 | 0% | 188 | 46% | 70 | 40% | 465 | 60% |
| Other<sup>c</sup> | 116 | 61% | 12 | 25% | 433 | 44% | 24 | 17% | 455 | 37% | 133 | 31% | 1,173 | 41% |
| Total | 1,617 | 67% | 143 | 24% | 8,891 | 67% | 103 | 30% | 6,627 | 36% | 1,248 | 44% | 18,629 | 54% |

<sup>a</sup>. % Rel= Percent Released
<sup>b</sup>. Includes: fox squirrel [*Sciurus niger*], northern flying squirrel [*Glaucomys sabrinus*], red squirrel [*Sciurus vulgaris*], and southern flying squirrel [*Glaucomys volans*].
<sup>c</sup>. Includes: orders Lagomorpha, Rodentia, and Eulipotyphla. Additional Lagomorpha species include: snowshoe hare [*Lepus americanus*]. Additional Eulipotyphla species include: eastern mole [*Scalopus aquaticus*], northern short-tailed shrew [*Blarina brevicauda*], and star-nosed mole [*Condylura cristata*]. Additional Rodentia species include: American beaver [*Castor canadensis*], black rat [*Rattus rattus*], brown rat [*Rattus norvegicus*], common vole [*Microtus arvalis*], deer mouse [*Peromyscus*] sp., eastern chipmunk [*Tamias striatus*], eastern woodrat [*Neotoma floridana*], house mouse [*Mus musculus*], meadow jumping Mouse [*Zapus hudsonius*], meadow vole [*Microtus pennsylvanicus*], muskrat [*Ondatra zibethicus*], North American porcupine [*Erethizon dorsatum*],

Rabies vector species (Table 8) were submitted less often (2,595), but had the second highest release rate (59.6%) of any group, behind the waterfowl. Although raccoons are common across New York State, only 7.3% (n = 1,856) of all mammals treated were raccoons; 402 skunks and 337 bats were reported by rehabilitators, together accounting for less than 3% of all mammals treated (n = 25,490). Particularly low numbers of admissions were recorded for the little brown bat [*Myotis lucifigus*] (n=46), which was once the most populous bat species in the state.

**Table 8.**
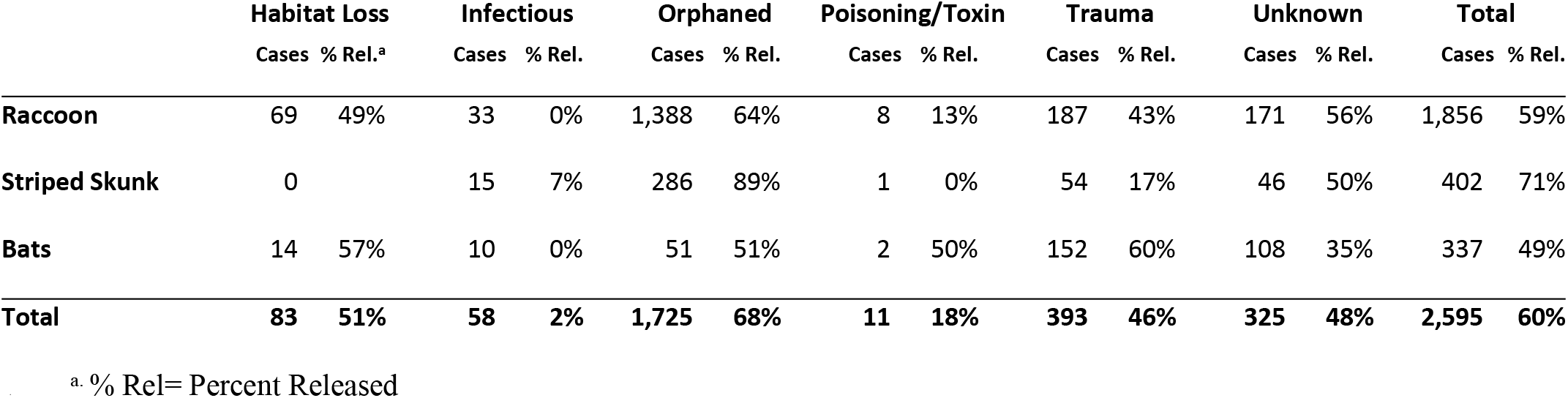
Cause of presentation and corresponding disposition for rabies vector species in rehabilitation.

The small carnivore group (Table 9) was comprised almost entirely of the Virginia opossum [*Didelphis virginiana*] (2,420). This species most frequently presented as orphaned with a 63.8% (n = 1,146/1,795) release rate. Overall, opossums had a release rate of 60.6% (n = 1,526/2,520).

**Table 9.** Cause of presentation and corresponding disposition for small carnivores in rehabilitation.

|  |  |  |  |  |  |  |  |  |  |  |  |  |  |  |
| --- | --- | --- | --- | --- | --- | --- | --- | --- | --- | --- | --- | --- | --- | --- |
| Virginia Opossum | 7 | 71% | 18 | 17% | 1,795 | 64% | 11 | 9% | 531 | 54% | 158 | 54% | 2,520 | 61% |
| Other<sup>b</sup> | 0 |  | 3 | 33% | 22 | 55% | 0 | 0% | 36 | 50% | 8 | 25% | 69 | 48% |
| **Total** | **7** | **71%** | **21** | **19%** | **1,817** | **64%** | **11** | **9%** | **567** | **53%** | **166** | **53%** | **2,589** | **60%** |

<sup>a</sup>. % Rel= Percent Released
<sup>b</sup>. Includes: American marten [*Martes americana*], American mink [*Neovison vison*], fisher [*Pekania pennanti*], least weasel [*Mustela nivalis*], long-tailed weasel [*Mustela frenata*], North American otter [*Lontra canadensis*], short-tailed weasel [*Mustela erminea*], and weasel [*Mustela*] sp.

Non-chelonian species were infrequently reported. Turtles presented most often for trauma, for which they had a release rate of 46.0%. Trauma due to collision with a car alone was responsible for 746 admissions, accounting for 32.7% of all admitted turtles. Rehabilitators also reported turtles as “orphans” 22.4% of the time, which had the lowest release rate for this distress category amongst the nine groups (n = 246/512, 48.0%) (Table 10).

**Table 10.**
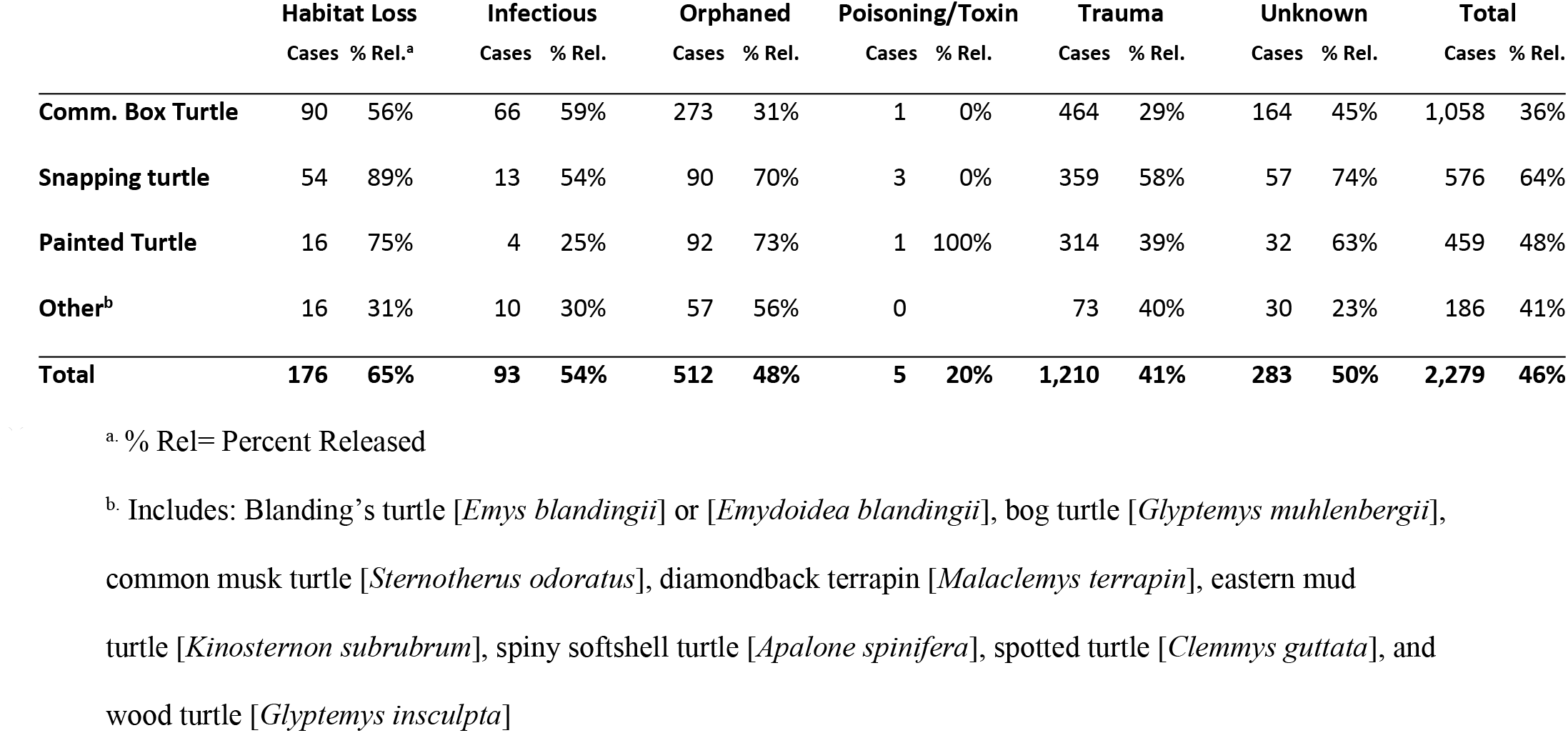
Cause of presentation and corresponding disposition for turtles in rehabilitation.

Box turtles comprised over 46% of all reptilian submissions, most of which presented as a result of trauma (43.9%). These turtles were successfully released 29% of the time. Young, “orphaned” box turtles were the next largest group, with a similar release rate of 31%. Vehicular-related injury accounted for 20% of all box turtle trauma with a release rate of 28%.

Known domestic cat attacks were responsible for 3,959 cases of wildlife rehabilitation during the study period. This was nearly twice the number of attacks by domestic dogs (2,068), and nearly 5 times the number of attacks by natural predators (837). Almost 5,000 animals were presented to rehabilitation centers with evidence of animal or human-inflicted wounds without proof of the inciting species. Mammals (2,002), mostly belonging to the small herbivore group (1,955), were the most frequent victims. Nearly 1,900 birds fell victim to the domestic cat, with 1,384 belonging to the passerine group. Wildlife known to have been injured by cats were successfully released 31% of the time, with 68% of victims dying as a result of their injuries; a loss of over 2,600 animals in a three-year span.

## Discussion

Previous wildlife rehabilitation literature has typically been limited to single species, diagnosis, or wildlife rehabilitation center of interest [14–17]. A strength of the current study and the DEC dataset is that the data encompass all cases across the entire state over a multiple year period, in a state that licenses rehabilitators to care for highly diverse species. Our results demonstrate the large community engaged in wildlife rehabilitation and the enormous scope of wildlife impacted by these practices. Key findings include significant increases in overall caseload and successful release since 1989, the concentration of over 40% of cases in just 2.3% of the licensed rehabilitators, identification of the most common causes of presentation/distress, and insight into potential population impacts for species of special concern. This information can provide a foundation for regulatory managemen, including where the state can focus rehabilitator educational efforts, determining allocation of resources benefiting specific species, and improving public outreach on anthropogenic impacts, such as free-roaming cats.

Although the traditional interactions between humans and wildlife in the form of hunting and trapping have declined, the rise in wildlife rehabilitation is demonstrates alternative ways the public engages with wildlife [18–20]. Wildlife rehabilitators are an avenue for the public to obtain information about native wildlife. Increasing awareness of defaunation and mass extinction [21,22] may motivate more individuals to “save” wildlife on a local level. In New York, the annual case numbers have increased substantially since 1989. Even the number of reptiles and amphibians under care has gone up, indicating a growing interest in this activity [13]. Increasing case numbers might reflect citizens’ increased awareness of wildlife species loss and willingness to engage with injured/orphaned wildlife. Alternatively, increasing case numbers might reflect higher human-wildlife interactions due to habitat fragmentation or increasingly distressed wildlife.

Agencies are tasked with managing the impacts and conflicts from wildlife rehabilitation, but often do not have specific resources to address issues [3,8,23]. However, the 1989 case data review and the present data indicate that nearly three-quarters of rehabilitators handled 25 or less cases per year, while less than 3% of rehabilitators handled more than 300 cases annually [13]. So within the rehabilitation community, a majority of the animals are cared for by a small number of centers; therefore significant data collection, communications, and outreach can be efficiently accomplished by targeting those resources.

The majority of animals presented to rehabilitators were impacted by trauma (38.2%) and orphaning (36.8%), much of which was likely anthropogenic. The proportion of injuries has increased since 1989 for both trauma (31.1%) and orphaning (40.2%). Over 1,000 animals each year were presented to NY rehabilitators as orphans due to confirmed unnecessary human intervention. These “orphaned” animals are generally healthy and are mistakenly taken for rehabilitation by well-meaning members of the public when they are in fact being cared for by a parent [24,25]. Improving public outreach regarding causes of wildlife injury and natural behaviors may help to reduce these numbers.

Probability of successful rehabilitation was dependent upon time in rehabilitation, species, and inciting cause. The overall release rate for rehabilitated animals was 50.2%, which was significantly higher than the reported rate in 1989. This rate may seem low, but should take into consideration the high standard of function necessary for a wild animal to be successful in the wild. Interestingly, eliminating the cohort of animals that died or were euthanized in the first 2 days raises the overall relase rate to 70.8%. Many of those animals lost in the first 48 hours are often the most severely injured or compromised. The resulting increased release rate for animals that survive the initial period supports a greater confidence in the quality and success of rehabilitative care.

Evaluation of release rates by cause of distress and species provides useful information to rehabilitators or veterinarians to determine the likelihood of successful rehabilitation for an individual case or allocate resources. Of all animals admitted due to orphaning, 65.7% were successfully released to the wild. In contrast, only 37.5% of trauma cases were successfully released to the wild with considerable species differences in release rates. For example, waterfowl as a group had the highest collective release rate (66.0%), including that for trauma victims (51.8%). For raptors, causes of distress documented by rehabilitators were consistent with previously published literature, which cited trauma as the most frequent cause of admission and mortality [26,27]. Raptors typically require high standards of function to be deemed releasable, with two perfectly functioning wings, intact talons, and unobscured bilateral vision for diurnal species, which may contribute to their lower release rates for traumatic injuries (40.4%). Data regarding release success for specific injuries (i.e. humeral fractures, ocular damage, shoulder luxation, etc.) are lacking and often lead to debate regarding the suitability of an animal for release. Collection of more detailed data regarding these injuries, treatments, and time spent in rehabilitation from these cases could be beneficial in further improving decision making.

For locally threatened or endangered species, rehabilitation of individuals may be of importance when the wild population is limited. Bald eagle populations have been closely monitored in New York since species restoration efforts began in 1976. In 2015, 264 breeding pairs were estimated statewide; consequently, 55 eagles seen in rehabilitation during this study period may represent a significant portion of the population [28]. The iconic nature of bald eagles makes them a prime candidate for citizens to submit to rehabilitation facilities if they are found injured or distressed. Rehabilitation of bald eagles is particularly challenging due to the species’ propensity for developing secondary complications. Bald eagles are highly susceptible to stress-related injuries, as well as aspergillosis-associated respiratory disease and staphylococcal pododermatitis [29–31]. Yet, the bald eagle had an overall release rate that was similar to other raptors, (45.5%), likely due to the signficant resources dedicated to their intake and care.

Bald eagles presented more frequently for poisoning/toxin than any other raptor species. This is likely indicative of lead toxicity, which has historically impacted the species heavily due to its scavenging feeding habits and position at the top of the food web [32]. Recent post-mortem studies estimate approximately 17% of bald eagles from New York State to have resulted from lead toxicity, and over 80% of all deceased eagles to have some exposure [33]. Nearly half of the bald eagles in our study population were presented to rehabilitation centers for an unknown cause. It is possible some of these admissions may have resulted from lead toxicity because the clinical signs can be vague, varying from malaise to profound neurologic consequences [34]. While lead analyzers are more frequently available in wildlife hospitals, they are expensive and specialized equipment not be readily available to most rehabilitators and veterinarians [35]. Toxicity-related presentations demonstrate potential large-scale health issues that may impacts species at the population level.

Wildlife rehabilitation data may also inform species health and natural history, as well as propose new avenues of research. Breeding pair estimates for waterfowl species in NYS, including wood ducks, are calculated via the Atlantic Flyway Northeast Plot Survey, which is carried out annually by the DEC. From 2012-2014, approximately 40 to 50,000 wood duck breeding pairs were estimated, making the 444 members of this species treated by wildlife rehabilitators representative of approximately 0.5% of the population [36]. Of note is the poor survival rate for the 374 orphaned ducklings that we report. Nesting holes are often constructed at heights up to 60 feet and ducklings leave the nest shortly after hatching [37]. The high incidence of orphaning amongst this species may be representative of premature fledging and/or trauma associated with a clumsy fall from a tall nest location, but more research into neonatal development and clutch survival rates is needed to determine more effective rehabilitation strategies for these ducklings and better understand wood duck natural history and population trends.

In New York, box turtles are listed as a species of special concern [38,39]. Population estimates are not available for the species in New York State, likely due to the difficulty of surverying elusive and solitary reptiles, however successful surveys have been performed in neighboring states [40]. Impacts of roadway construction and an increasing density of vehicular traffic were exemplified in the large proportion of turtles presented following car collision. While box turtles represented the most commonly presented reptile following vehicular trauma, this was in lesser proportion than the common snapping turtle (*Chelydra serpentine*) and painted turtle (*Chyrsemys picta*). Overall, however, the aquatic turtles were successfully rehabilitated following vehicular trauma more frequently than the box turtle, for reasons that are unclear. The species’ poor release rate emphasizes the importance of motorist education and habitat protection for this state-recognized species if we are to combat its precipitous decline.

Wildlife rehabilitation data may help to address controversial topics in ecologic health and public policy. The impact of free-roaming domestic cats on wildlife populations has been heavily debated [1,14,15,24,25,41]. We demonstrated nearly 4,000 animals known to be injured by cats over three years, with an additional 5,000 animals with wounds possibly inflicted by a predator. Known domestic cat attacks accounted for 6.6% of all wildlife rehabilitation cases in New York State. Previous studies have reported up to 14% of all admissions to rehabilitation centers to be a result of domestic pet interaction [25]. Less than one-third of cat-attack victims were successfully released, which closely mirrored prior literature describing the release rates for birds attacked by cats to be as low as 12.9% [24]. Additionally, species less commonly considered to be victims of cats, including lizards and bats, have been documented to suffer substantially from feline predation [14,15]. In addition to birds and small mammals, our study documented several species of bats, turtles, and snakes to be confirmed victims of domestic cat predation. Quantitative evidence for the debilitating impact that free-ranging domestic cats have on wildlife may be more impactful in public education and policy regulation.

Since wildlife rehabilitators handle large numbers of animals, we are able to track trends in species over time as an indicator of disease activity or population health. The American crow, known to suffer high mortality from West Nile Virus (WNV) [42–45], had the lowest overall release rate of any species. Real-time case reporting since the introduction of WNV in 1999 would have detected increased crow mortality, while annual case tracking combined with confirmatory testing could help elucidate ecological factors impacting WNV prevalence. Similarly, the small number of bats rehabilitated likely reflects the emerging fungal disease known as White Nose Syndrome (WNS). Previously New York’s most common bat species, the little brown bat population has been devastated by WNS since its arrival in 2006; the population has experienced a 90% decrease and has the potential for extinction by 2026 [46,47]. Submissions to the state’s rabies laboratory also show a precipitous decline in this species over the same time period [48].

Trained rehabilitators serve as an important buffer between the public and wildlife disease threats. When rehabilitation is restricted or prohibited, members of the public may be reticent to abandon wild animals or submit them for immediate euthanasia. These animals may then be raised in inappropriate conditions, increasing to possibility of zoonotic disease transmission. Public health agencies can be tasked with animal confiscation while conducting expensive and time consuming follow-up to control zoonotic disease exposures, such as rabies. In New York, additional requirements are necessary to obtain a license to rehabilitate rabies vector species. These include a minimum of two years experience as a rehabilitator, pre-exposure rabies vaccinations, attendance at an annual training session, additional record-keeping responsibilities, and an inspection of the rehabilitation facility by the state’s Department of Agriculture and Markets [49]. Over 2,500 raccoons, skunks, and bats were submitted for rehabilitation during the reporting period so the potential exposure to the public would be substantial if trained rehabilitators were not available.

We recognize several limitations to the current study. While the records represent all wildlife rehabilitation submissions over a three-year span, reporting errors and inconsistencies both within and between rehabilitation centers occurred. These ranged from spelling errors, to misidentified species, to failure to accurately report distress causes and final disposition. While selecting the primary distress for each case was done systematically for every possible combination, the process represents potential for bias from the authors. Finally, we define rehabilitation success as release to the wild. While true success may be defined as normal function and survival after the time of release, this information is unfortunately not available for most species on a large scale post-rehabilitation. The lack of case follow-up is an inherent shortcoming in the current process of wildlife rehabilitation.

Wildlife rehabilitation plays an important role in public education and outreach about native wildlife. There are benefits for animal welfare, disease monitoring, and conservation [50–53]. Analysis of large-scale rehabilitation data can improve resource allocation, treatment methods, surveillance, public education, and regulatory decision making. Standardizing rehabilitator reporting and digital data collection would facilitate compilation and analysis to benefit both rehabilitators and the state agency regulators.

## Acknowledgements

The authors would like to thank the New York State Department of Environmental Conservation, the New York State Wildlife Rehabilitation Council, and Cornell University students Lily Cheng, Richalice Melendez, and Mariacamila Garcia Estrella who contributed to data compilation and analysis. Funding support was provided by the New York State Department of Environmental Conservation through the Federal Aid in Wildlife Restoration.

## References

1. Schenk AN, Souza MJ. Major Anthropogenic Causes for and Outcomes of Wild Animal Presentation to a Wildlife Clinic in East Tennessee, USA, 2000–2011. PLoS One. 2014;9. doi:10.1371/journal.pone.0093517

2. Vogelnest L, Woods R. Medicine of Australian Mammals. 2008.

3. Deem SL, Karesh WB, Weisman W. Putting Theory into Practice: Wildlife Health in Conservation. Conserv Biol. 2001;15: 1224–1233.

4. Moore M, Early G, Touhey K, Gulland F, Wells R. Rehabilitation and Release of Marine Mammals in the United States: Risks and Benefits. Mar Mammal Sci. 2007;23: 731–750. doi:10.1111/j.1748-7692.2007.00146.x

5. Mullineaux E. Veterinary treatment and rehabilitation of indigenous wildlife. J Small Anim Pract. 2014;55: 293–300. doi:10.1111/jsap.12213

6. Siemer WF, Brown TL. Wildlife Rehabilitators’ Attitudes and Motivations: Insights from New York. Ithaca, NY; 1992. pp. 205–220;.

7. National Wildlife Rehabilitators Association. Our Members. 2015.

8. Stitt T, Mountifield J, Stephen C. Opportunities and obstacles to collecting wildlife disease data for public health purposes: Results of a pilot study on Vancouver Island, British Columbia. 2011;48.

9. Orós J, Montesdeoca N, Camacho M, Arencibia A, Calabuig P. Causes of Stranding and Mortality, and Final Disposition of Loggerhead Sea Turtles (Caretta caretta) Admitted to a Wildlife Rehabilitation Center in Gran Canaria Island, Spain (1998-2014): A Long-Term Retrospective Study. Ambrósio CE, editor. PLoS One. 2016;11: 1–14. doi:10.1371/journal.pone.0149398

10. New York State Department of Environmental Conservation Division of Fish and Wildlife. Checklist of Amphibians, Reptiles, Birds and Mammals of New York State Including Their Legal Status. Albany; 2019.

11. The New York State Ornithological Association I. Checklist of the Birds of New York State. 2019.

12. Loss SR, Will T, Marra PP. The impact of free-ranging domestic cats on wildlife of the United States. Nat Commun. 2013.

13. Siemer WF, Brown TL. Characteristics, activities, and attitudes of licensed wildlife rehabilitators in New York. Ithaca, NY; 1992.

14. Koenig J, Shine R, Shea G. The Dangers of Life in the City: Patterns of Activity, Injury and Mortality in Suburban Lizards (Tiliqua scincoides). J Herpetol. 2002;36: 62–68. doi:10.1670/0022-1511(2002)036[0062:TDOLIT]2.0.CO;2

15. Ancillotto L, Tiziana Serangeli M, Russo D. Curiosity killed the bat: Domestic cats as bat predators. Mamm Biol. 2013;78: 369–373. doi:10.1016/j.mambio.2013.01.003

16. Burton E, Tribe A. The rescue and rehabilitation of koalas (Phascolarctos cinereus) in Southeast Queensland. Animals. 2016;6: 2–10. doi:10.3390/ani6090056

17. Page-Karjian A, Norton TM, Krimer P, Groner M, Nelson SE, Gottdenker NL. Factors Influencing Survivorship of Rehabilitating Green Sea Turtles (Chelonia mydas) with Fibropapillomatosis. J Zoo Wildl Med. 2014;45: 507–519. doi:10.1638/2013-0132R1.1

18. Matheny K. Michigan Hunting in Major Decline - Why That Matters. Detroit Free Press. 10 Nov 2018.

19. U.S. Department of the Interior, U.S. Fish and Wildlife Service, and U.S. Department of Commerce USCB. 2016 National Survey of Fishing, Hunting, and Wildlife-Associated Recreation. 2016.

20. Decker D, Smith C, Forstchen A, Hare D, Pomeranz E, Doyle-capitman C, et al. Governance Principles for Wildlife Conservation in the 21st Century. 2016;9: 290–295. doi:10.1111/conl.12211

21. Ceballos G, Ehrlich PR, Dirzo R. Biological annihilation via the ongoing sixth mass extinction signaled by vertebrate population losses and declines. Proc Natl Acad Sci U S A. 2017;114: E6089–E6096. doi:10.1073/pnas.1704949114

22. Dirzo R, Young HS, Galetti M, Ceballos G, Isaac NJ, Collen B. Defaunation in the Anthropocene. Sci Mag. 2014;345: 401–406.

23. Barnes E. To what extent are veterinary practices prepared to treat wildlife patients? A cross-sectional study of perceptions of responsibility and capability of treating wildlife in UK veterinary practices. Plymouth Student Sci. 2017;10: 1–21.

24. Mcruer DL, Gray LC, Horne LA, Clark EE. Free-roaming cat interactions with wildlife admitted to a wildlife hospital. Journal of Wildlife Management. 2017. pp. 163–173;. doi:10.1002/jwmg.21181

25. Loyd KAT, Hernandez SM, McRuer DL. The role of domestic cats in the admission of injured wildlife at rehabilitation and rescue centers. Wildl Soc Bull. 2017;41: 55–61. doi:10.1002/wsb.737

26. Montesdeoca N, Calabuig P, Corbera JA, Orós J. Causes of Admission for Raptors to the Tafira Wildlife Rehabilitation Center, Gran Canaria Island, Spain: 2003-13. J Wildl Dis. 2016;52: 647–652. doi:10.7589/2015-09-255

27. Smith KA, Campbell GD, Pearl DL, Jardine CM, Salgado-Bierman F, Nemeth NM. A Retrospective Summary of Raptor Mortality in Ontario, Canada (1991-2014), Including the Effects of West Nile Virus. J Wildl Dis. 2017;54: 2017-07–157. doi:10.7589/2017-07-157

28. New York State Department of Environmental Conservation. Conservation plan for bald eagles in New York State. 2016.

29. Deem SL. Fungal diseases of birds of prey. Vet Clin North Am Exot Anim Pract. 2003;6: 363–376. doi:10.1016/S1094-9194(03)00004-5

30. Harris MC, Sleeman JM. Morbidity and mortality of bald eagles (Haliaeetus leucocephalus) and peregrine falcons (Falco peregrinus) admitted to the Wildlife Center of Virginia, 1993-2003. J Zoo Wildl Med. 2007;38: 62–66. doi:10.1638/05-099.1

31. Rodriguez-Lainz AJ, Hird DW, Kass PH, Brooks DL. Incidence and risk factors for bumblefoot (pododermatitis) in rehabilitated raptors. Prev Vet Med. 1997;31: 175–184. doi:10.1016/S0167-5877(96)01137-3

32. Martin PA, Hughes · K D, Campbell · G D, Shutt · J L. Metals and Organohalogen Contaminants in Bald Eagles (Haliaeetus leucocephalus) from Ontario, 1991–2008. Arch Environ Contam Toxicol. 2017;1. doi:10.1007/s00244-017-0479-5

33. Cornell Wildlife Health Lab. Bald Eagles and Lead Toxicity 2017. 2017 [cited 17 Oct 2019]. Available: https://cwhl.vet.cornell.edu/project/bald-eagles-and-lead-toxicity-2017

34. De Francisco N, Ruiz Troya J, Agüera E, Aguëra E. Lead and lead toxicity in domestic and free living birds. Avian Pathol. 2010;32: 3–13. doi:10.1080/0307945021000070660

35. Herring G, Eagles-Smith CA, Bedrosian B, Craighead D, Domenech R, Langner HW, et al. Critically assessing the utility of portable lead analyzers for wildlife conservation. Wildl Soc Bull. 2018;42: 284–294. doi:10.1002/wsb.892

36. Migratory Game Bird Management and Surveys. In: NYS Dept. of Environmental Conservation [Internet]. [cited 29 Oct 2019]. Available: https://www.dec.ny.gov/outdoor/115095.html

37. Cornell Lab of Ornithology. Wood Duck Life History. In: All About Birds [Internet]. [cited 18 Oct 2019]. Available: https://www.allaboutbirds.org/guide/Wood_Duck/lifehistory

38. NYS Dept. of Environmental Conservation. List of Endangered, Threatened and Special Concern Fish & Wildlife Species of New York State. 2015 [cited 18 Oct 2019]. Available: https://www.dec.ny.gov/animals/7494.html

39. Rules and Regulations of the State of New York. 182. 4 Listing of species of special concern. 2019.

40. Erb LA, Willey LL, Johnson LM, Hines JE, Cook RP. Detecting long-term population trends for an elusive reptile species. J Wildl Manage. 2015;79: 1062–1071. doi:10.1002/jwmg.921

41. Loss SR, Will T, Marra PP. The impact of free-ranging domestic cats on wildlife of the United States. Nat Commun. 2013;4: 1396. doi:10.1038/ncomms2380

42. Nemeth NM, Beckett S, Edwards E, Klenk K, Komar N. Avian mortality surveillance for West Nile virus in Colorado. Am J Trop Med Hyg. 2007;76: 431–437. doi:76/3/431 [pii]

43. Eidson M, Kramer L, Stone W, Hagiwara Y, Schmit K. Dead Bird Surveillance as an Early Warning System for West Nile Virus. Emerg Infect Dis. 2001;7.

44. Eidson M, Komar N, Sorhage F, Nelson R, Talbot T, Mostashari F, et al. Crow deaths as a sentinel surveillance system for West Nile virus in the northeastern United States, 1999. Emerg Infect Dis. 2001;7: 615–20. doi:10.3201/eid0704.010402

45. Bernard KA, Maffei JG, Jones SA, Kauffman EB, Ebel GD, Dupuis AP, et al. West Nile Virus Infection in Birds and Mosquitoes, New York State, 2000. Emerg Infect Dis. 2001;7: 679–685. doi:10.3201/eid0704.017415

46. Frick WF, Pollock JF, Hicks AC, Langwig KE, Reynolds DS, Turner GG, et al. An emerging disease causes regional population collapse of a common North American bat species. Science. 2010;329: 679–82. doi:10.1126/science.1188594

47. Blehert DS, Hicks AC, Behr M, Meteyer CU, Berlowski-Zier BM, Buckles EL, et al. Bat White-Nose Syndrome: An Emerging Fungal Pathogen? doi:10.1126/science.1163874

48. Rabies Laboratory Wadsworth Center NYSD of H. 2015 Rabies Annual Report. Albany; 2015.

49. New York State Department of Environmental Conservation. Wildlife Rehabilitator License - NYS Dept. of Environmental Conservation. [cited 12 Feb 2018]. Available: https://www.dec.ny.gov/permits/25027.html

50. Ana A, Perez Andrés M, Julia P, Pedro P, Arno W, Kimberly VW, et al. Syndromic surveillance for West Nile virus using raptors in rehabilitation. BMC Vet Res. 2017;13. doi:10.1186/s12917-017-1292-0

51. Tribe A, Brown PR. The role of wildlife rescue groups in the care and rehabilitation of Australian fauna. Hum Dimens Wildl. 2000;5: 69–85. doi:10.1080/10871200009359180

52. Rio-Maior H, Beja P, Nakamura M, Santos N, Brandão R, Sargo R, et al. Rehabilitation and post-release monitoring of two wolves with severe injuries. J Wildl Manage. 2016;80: 729–735. doi:10.1002/jwmg.1055

53. Baker L, Edwards W, Pike D. Sea turtle rehabilitation success increases with body size and differs among species. Endanger Species Res. 2015;29: 13–21. doi:10.3354/esr00696

